# Comparative Transcriptome and Methylome Analysis in Human Skeletal Muscle Anabolism, Hypertrophy and Epigenetic Memory

**DOI:** 10.1101/465708

**Authors:** Daniel C. Turner, Robert A. Seaborne, Adam P. Sharples

## Abstract

Transcriptome wide changes in human skeletal muscle after acute (anabolic) and chronic resistance exercise (RE) induced hypertrophy have been extensively determined in the literature. We have also recently undertaken DNA methylome analysis (850,000 + CpG sites) in human skeletal muscle after acute and chronic RE, detaining and retraining, where we identified an association between DNA methylation in an epigenetic memory of exercise induced skeletal muscle hypertrophy. However, it is currently unknown as to whether all the genes identified in the transcriptome studies to date are also epigenetically regulated at the DNA level after acute, chronic or repeated RE exposure. We therefore aimed to undertake large scale bioinformatical analysis by pooling the publicly available transcriptome data after acute (110 samples) and chronic RE (181 samples) and comparing these large data sets with our genome-wide DNA methylation analysis in human skeletal muscle after acute and chronic RE, detraining and retraining. Indeed, after acute RE we identified 866 up- and 936 down-regulated genes at the expression level, with 270 (out of the 866 up- regulated) identified as being hypomethylated, and 216 (out of 936 downregulated) as hypermethylated. After chronic RE we identified 2,018 up- and 430 down-regulated genes with 592 (out of 2,018 upregulated) identified as being hypomethylated and 98 (out of 430 genes downregulated) as hypermethylated. After KEGG pathway analysis, genes associated with ‘cancer’ pathways were significantly enriched in both bioinformatic analysis of the pooled transcriptome and methylome data after both acute and chronic RE. This resulted in 23 (out of 69) and 28 (out of 49) upregulated and hypomethylated and 12 (out of 37) and 2 (out of 4) downregulated and hypermethylated ‘cancer’ genes following acute and chronic RE respectively. Within skeletal muscle tissue, these ‘cancer’ genes predominant functions were associated with matrix/ actin structure and remodelling, mechano-transduction (including PTK2/Focal Adhesion Kinase and Phospholipase D-following chronic RE only), TGF-beta signalling and protein synthesis (GSK3B after acute RE only). Interestingly, 51 genes were also identified to be up/downregulated in both the acute and chronic RE pooled transcriptome analysis as well as significantly hypo/hypermethylated after acute RE, chronic RE, detraining and retraining. Five genes; FLNB, MYH9, SRGAP1, SRGN, ZMIZ1 demonstrated increased gene expression in the acute and chronic RE transcriptome and also demonstrated hypomethylation in these conditions. Importantly, these 5 genes demonstrated retained hypomethylation even during detraining (following training induced hypertrophy) when exercise was ceased and lean mass returned to baseline (pre-training) levels, identifying them as potential epigenetic memory genes. Importantly, for the first time across the transcriptome and epigenome combined, this study identifies novel differentially methylated genes associated with human skeletal muscle anabolism, hypertrophy and epigenetic memory.

## Introduction

Skeletal muscle tissue demonstrates extensive plasticity, responding dynamically to sustained mechanical loading and contraction with muscle hypertrophy. However, skeletal muscle tissue also wastes (atrophy) rapidly during periods of disuse, for example, following an injury from a fall or reduces in size overtime as a result of ageing (sarcopenia, reviewed in ^1, 2^). The transcriptome wide changes in gene expression that regulate healthy adult human skeletal muscle anabolism and hypertrophy in response to acute and chronic resistance exercise (RE) respectively have been reported in the literature ^3–10^. Ultimately, the identification of genes associated with skeletal muscle mass regulation continue to progress this field of research in order to optimise the growth response to resistance exercise and help prevent muscle wasting. Despite these recent advances, it is currently unknown as to whether the genes identified at the mRNA level across the transcriptome are also epigenetically regulated at the DNA level.

Epigenetics is the study of DNA that is modified as a result of an encounter with the environment. These DNA modifications subsequently affect genes at the transcript level. The major forms of DNA modification include alterations to the surrounding histones as a result of methylation, acetylation and deacetylation. Histone modifications lead to the DNA being rendered into a repressive (inhibitory) or permissive (allowing) state, that subsequently alters access for the transcriptional machinery regulating gene expression. DNA itself can also be modified directly by methylation, via the addition or removal of methyl groups, particularly to cytosine-guanine base pairing (CpG) sites. For example, increased DNA methylation (hypermethylation) that occurs in the fifth position of a cytosine (5mC) residue of a CpG site, particularly if present in the promoter or enhancer region of a gene, can attenuate the performance of transcriptional apparatus and can cause a reduction in the expression of a specific gene ^11^. On the other hand, a reduction in DNA methylation (hypomethylation) can improve the function of transcription factors and therefore enhance gene expression ^11^.

We have recently characterised genome-wide DNA methylation of over 850,000 CpG sites across the human genome in skeletal muscle at rest (baseline), after acute resistance exercise (acute RE), chronic resistance exercise induced hypertrophy (training/loading), followed by a period of detaining (where muscle returned to baseline- termed unloading), and finally after retraining induced hypertrophy (retraining/reloading)^12^. Therefore, in the present study we aimed to identify if transcriptome wide gene expression changes were associated with DNA methylation of the same genes across the genome (methylome) after acute and chronic RE. In order to achieve this aim, we undertook large scale bioinformatical analysis that pooled the publicly available transcriptome data sets from human skeletal muscle pre/post-acute (110 gene array samples) and chronic (181 gene array samples) resistance exercise and overlapped this with the genome-wide DNA methylation after acute and chronic resistance exercise from our recent study ^12^.

Finally, following exciting data that defined a cellular memory in skeletal muscle ^13, 14^. we have recently demonstrated that skeletal muscle has an epigenetic memory ^12, 15^. Where, DNA methylation signatures (particularly hypomethylation) were retained even when exercise ceased during detraining (and lean mass was returned to pre-exercise levels) following chronic RE induced hypertrophy in humans, leading to an advantageous state for these genes to be further upregulated when later retaining induced hypertrophy was encountered ^12^. However, in this study we did not perform transcriptome wide gene expression analyses, opting instead for targeted gene expression analysis of those genes that mapped to the most significantly regulated CpG sites. Therefore, the secondary aim of the present study was to identify if transcriptome wide gene expression changes from pooled publicly available data sets after acute and chronic RE, were associated with the methylome changes seen during training, detraining and retaining in our epigenetic muscle memory studies.

Overall, we hypothesised that we would identify new epigenetically regulated genes and gene pathways in human skeletal muscle anabolism, hypertrophy and those associated with epigenetic muscle memory. We also hypothesised that transcriptome wide changes identified in the pooled transcriptome analysis after acute and chronic RE would be associated with altered DNA methylation of some of the same genes after training, detraining and retraining.

## Methods

### Identification of transcriptomic studies

The studies used for pooled transcriptome analysis for both acute and chronic resistance exercise (RE) are summarised in Suppl. File 1. These include publicly available transcriptome data sets deposited and searchable before the end of April 2018. In order to enable a suitable comparison of the transcriptomic studies after acute and chronic RE in humans we incorporated the following inclusion and exclusion criteria in our search. For acute RE, we included all transcriptome array data from the skeletal muscle of young healthy adult males in order to enable the comparison with the genome-wide DNA methylation analysis that was undertaken in the same population in ^12^. Resistance exercise intensity was not an exclusion criterion per se, however no aerobic, continuous or concurrent exercise studies were included. All time points of isolated RNA from biopsies that were within the first 24 hrs post an acute bout of RE were included, and samples taken after 24 hrs were excluded. Due to the varied time points at which RNA was sampled following acute RE, with only a small number of studies sampling at the same timepoint, all samples taken immediately post exercise and up to 24 hrs were attributed to the same post-acute RE biopsy condition for the pooled transcriptome analysis. Furthermore, within studies, if there were multiple time points of biopsy sampling, all times points were included up to 24 hrs and attributed within the same post-acute RE biopsy condition for the analysis. We excluded any study with less than 10,000 gene probe-sets annotated by ‘gene symbol’. These studies used earlier gene array platforms with a low number of gene transcripts analysed versus more recent data sets with approximately 20,000 or more gene transcripts. When including the studies with 10,000 probe-sets or less, this dramatically affected the number of the same genes that were assessed between studies and meant that we could not accurately compare any of the larger more recent studies that used the latest gene array platforms. For chronic RE transcriptome data, we included young heathy adult males both pre and post chronic RE with biopsies taken at rest. There was no set frequency, duration or intensity of RE-although minimum frequency and duration for the chronic RE studies was 12 weeks resistance exercise 2 times per week (detailed in Suppl. File 1). As in the acute analysis above, we excluded probe sets below 10,000 annotated genes by ‘gene symbol.’ A table of all studies can be found in Suppl. File 1.

### Bioinformatic Pooled Transcriptome Analysis

All available data sets were downloaded from NIH data base, *Gene Expression Omnibus* (GEO), website and imported into Partek Genomics Suite software (version 7.18.0518, Partek Inc. Missouri, USA) as .CEL or .TXT files for both acute and chronic RE studies independently. The relevant gene array platform annotation file, also downloaded from GEO, was then assigned to the appropriate data set. If intensity values were not Log transformed, these were subsequently Log transformed prior to continuing. For genes that were detected by more than one probe set we selected the highest-scoring probe set to represent that gene in order to allow a comparison of the unambiguous expression estimate of an individual gene, and compare it across studies/array platforms. This has previously been defined as a relevant method to account for altered hybridisation efficiency across multiple probe sets for the same gene and across array platforms ^16, 17^. To be able to directly compare gene expression between studies, we then identified a list of the common gene symbol annotations across all studies. This allowed us to filter each individual data set by gene symbols that were uniformly shared across the different array platforms, subsequently allowing comparison of identical genes across all included studies. All filtered acute RE or chronic RE data sets were then merged together and samples that did not satisfy the inclusion/exclusion criteria described above, were removed. Sample attributes were then defined as ‘pre’ or ‘post’ acute RE and ‘pre’ or ‘post’ chronic RE, also in line with the inclusion/exclusion criteria stated above. Quality assurance (QA) and control (QC) analyses were undertaken on both the acute and chronic study data sets independently. Principal component analysis, box/whisker charts and sample frequency/density plots by lines were produced for both the merged acute RE and chronic RE data sets. Outlier samples were detected using principle component analysis (PCA) and analysing the normal distribution of ß-values. Outliers were then removed if they fell outside 2 standard deviations (SDs) from the centroid using ellipsoids (Suppl. File 1 & 2) as well as showing different distribution patterns to the samples of the same condition (pre/post) or within each study. To correct for differences in processing methods of the samples across studies, we performed batch correction using Partek Genomics Suite ‘Remove Batch Effects’ tool (uses ANOVA approach), previously reported as an appropriate batch correction tool when dealing with data of non-trivial size (i.e. large number of samples per condition ^18, 19^). Specific and detailed description of QA/QC and batch removal can be seen in the appropriate figure legends. Detection of differentially expressed genes was then performed on Partek using an ANOVA for both acute and chronic RE (Pre vs. Post contrast) (version 7.18.0518) and a gene list of all the significantly (P ≤ 0.01) differentially regulated genes (both up and down) was created.

### Overlapping of Pooled Transcriptome Analysis with the DNA Methylome

Using Venn diagram analysis, the significant differentially regulated gene lists generated from the pooled transcriptomic gene expression analysis described above were overlapped with the significantly differentially modified CpG sites lists from the freely available supplementary files of ^12^, after both acute RE and chronic RE (latter termed ‘loading’ in the original article), detraining (unloading) and retaining (reloading). Genome-wide DNA methylation data were therefore processed as previously defined in the original data ^12^ and a recently published data descriptor for the original study ^20^. Briefly, raw .IDAT files were processed on Partek Genomic Suite V.6.6 (Partek Inc. Missouri, USA) and background normalisation was performed via the Subset-Quantile Within Array Normalisation (SWAN) method ^21^ and imported using the MethylationEPIC_v-1-0_B2.bpm manifest file. This analysis enabled us to compare genes that were both up/downregulated at the expression level and hypo/hypermethylated at the DNA level after both acute and chronic RE. It also allowed us to associate genes that were up/down regulated in the comparative transcriptome analysis after acute and chronic RE that were hypo/hypermethylated at the DNA level after acute RE, training, detaining and retaining.

### Pathway Analysis

Using statistically generated gene expression and CpG data (detailed above), KEGG signalling pathway analysis ^22–24^ (see results/figures for significance level of enrichment P values) was performed in Partek Genomic Suite and Partek Pathway. Once significant enrichment was determined in a Venn diagram analysis we overlapped the significant pathways common to both the transcriptome and methylome analysis.

### Target Gene Expression of FLNB by rt-qRT-PCR

Human skeletal muscle RNA previously derived in ^12^ was reanalysed and used in the present study. The RNA was originally isolated from biopsies taken from the vastus lateralis of young adult males. Participant characteristics, full biopsy and RNA isolation procedures are described in ^12, 20^. RNA from baseline, 30 minutes post-acute resistance exercise (acute RE), 7 weeks training (loading), 7 weeks detraining (unloading) and 7 weeks retraining (reloading) was used for rt-qRT-PCR using QuantiFastTM SYBR^®^ Green RT-PCR one-step kit on a Rotorgene 3000Q, with a reaction setup as follows; 4.75 μl experimental sample (7.36 ng/μl totalling 35 ng per reaction), 0.075 μl of both forward and reverse primer of the gene of interest (100 μM stock suspension), 0.1 μl of QuantiFast RT Mix (Qiagen, Manchester, UK) and 5 μl of QuantiFast SYBR Green RT-PCR Master Mix (Qiagen, Manchester, UK). Reverse transcription initiated with a hold at 50°C for 10 minutes (cDNA synthesis) and a 5 minute hold at 95°C (transcriptase inactivation and initial denaturation), before 40-45 PCR cycles of; 95°C for 10 sec (denaturation) followed by 60°C for 30 secs (annealing and extension). Primer sequence for gene of interest, Filamin B (FLNB), was designed to detect/span all FLNB transcript variants 1, 2, 3, 4 and X1 (NM_001164317.1, NM_001457.3, NM_001164318.1, NM_001164319.1 and XM_005264978.2 respectively) as a global measure of FLNB gene expression. Primer sequences for the gene of interest, FLNB were: F: GTGGACACCAGCAGGATCAA; R: CGGCCGAGAGTCAACTGTAA, product length 96 bp) and reference gene, RPL13a (NM_012423) were: F: GGCTAAACAGGTACTGCTGGG; R: AGGAAAGCCAGGTACTTCAACTT, product length 105 bp. Both FLNB and RPL13a primer sequences demonstrated no unintended targets via BLAST search and yielded a single peak after melt curve analysis conducted after the PCR step above. All relative gene expression was quantified using the comparative Ct (^∆∆^Ct) method ^25^. Individual participants own baseline (pre) Ct values were used in ^∆∆^Ct equation as the calibrator, using RPL13a as the reference gene, except for one of the participants where there was no baseline/pre RNA remaining from the original study^12^. Therefore, a pooled baseline/pre-value of all participants was used to generate fold changes in all other experimental conditions for this particular participant. Indeed, the pooled baseline samples for FLNB across all participants demonstrated a small variation in Ct value 25.59 (± 0.87 SDEV, 3.4% variation). The average Ct value for the reference gene RPL13a was also consistent across all participants and experimental conditions (22.04 ± 0.52, SDEV) with small variation of 2.3%. The average PCR efficiencies of FLNB (95.83%) were comparable with the reference gene RPL13a (97.07%).

## Results

### Pooled Transcriptome Analysis overlapped with the Methylome in Human Skeletal Muscle after Acute Resistance Exercise

In total, 110 gene arrays from skeletal muscle biopsies were used for pooled transcriptome analysis, including: 41 baseline (pre) samples and 69 post-acute resistance exercise (RE) samples (37 pre / 57 post after outlier removal). These studies shared 14,992 annotated genes by ‘gene symbol’ (Suppl. Figure 1A). Relevant QA/QC and outlier removal is depicted and described in Suppl. Figure 1B-I and detailed in the associated figure legend. Downstream differential gene expression analysis of the transcriptome data across all 5 studies demonstrated that there were 1,802 genes significantly differentially regulated (P ≤ 0.01) after acute RE (Full List Suppl. File 2A). Out of these genes, 866 were upregulated and 936 gene downregulated. Out of the 866 genes upregulated, 270 of these genes were also hypomethylated (Figure 1A, full gene list Suppl. File 2B), compared with 216 that demonstrated upregulation of gene expression and hypermethylation. Of the genes upregulated and hypomethylated (more likely to affect transcription), this equated to 355 different CpG sites on these 270 genes as some individual genes had more than one CpG site that was hypomethylated per gene (Suppl. File 2C). Furthermore, out of 936 downregulated genes there were 216 genes across pooled transcriptome studies that were also hypermethylated in the methylome analysis, Figure 1B, Suppl. File 2D), compared with 298 genes that were downregulated and hypomethylated. For the downregulated and hypermethylated genes (more likely to affect transcription) this equated to 268 CpG sites on these 216 genes (Suppl. File 2E).

**Figure 1:**
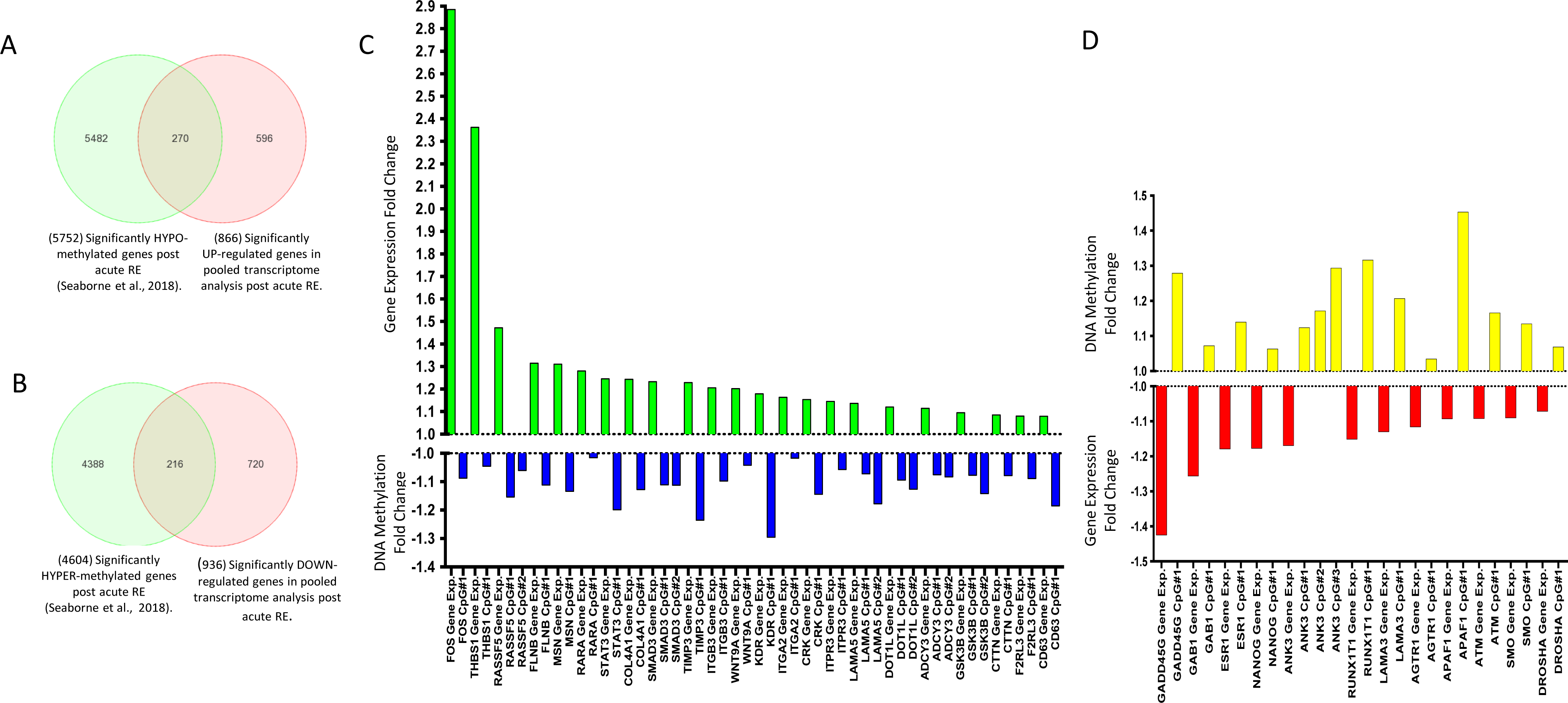
**A**, Venn Diagram demonstrating out of 866 genes upregulated after acute RE (p ≤ 0.01) in the pooled transcriptomic analysis, 270 of these genes were significantly hypomethylated in the methylome analysis ^12^. **B**, Venn Diagram demonstrating out of 936 genes downregulated after acute RE (p ≤ 0.01) in the pooled transcriptomic analysis, 216 of these genes were significantly hypermethylated in the methylome analysis ^12^. Note: 5752 / 4604 total hypo/hypermethylated CpGs after acute RE is different than the original paper reporting 9,153 / 8,212 hypo/hypermethylated CpGs. This is due to the number of CpG’s that resided on the shared list of 14,992 annotated genes by ‘gene symbol’ across the pooled transcriptomic studies for acute RE, depicted above in **1A**, in order to provide a direct comparison of CpG sites on the same genes from the pooled transcriptomic data set. **C.** ‘Cancer’ pathway genes upregulated (GREEN bars) in pooled transcriptomic studies (p ≤ 0.01) and hypomethylated (BLUE bars) in ^12^ (p ≤ 0.05) after acute RE. Fold change is Log transformed. In skeletal muscle, the majority of these genes (13 out of 23) are associated with matrix / actin structure or remodelling and mechano-transduction in skeletal muscle (MSN THBS1, TIMP3, FLNB, LAMA5, CRK, COL4A1, ITGA2, ITGB3, CD63, CTTN, RASSF5, F2RL3) and 3 genes with TGF-Beta signalling (SMAD3, FOS, WNT9A), 2 genes with calcium signalling (ITPR3, ADCY3), 1 gene with IL-6 signalling (STAT3), and protein synthesis (GSK3B) and retinoic acid signalling (RARA). **D**, ‘Cancer’ genes downregulated (RED) in pooled transcriptomic studies and hypermethylated (YELLOW bars) in after acute RE ^12^.

### Cancer Pathways are Enriched in both the Transcriptome and Methylome Data Sets in Human Skeletal Muscle after Acute Resistance Exercise

KEGG Pathway analysis of the top 20 pathways included 464 genes that were significantly differentially regulated (P ≤ 0.01) and significantly enriched (Enrichment P value ≤ 0.003). Out of these 468 genes, 36% (168 genes) were in 5 different enriched ‘cancer’ pathways (including; ‘Proteoglycans in cancer’, ‘Transcriptional misregulation in cancer’, ‘Colorectal cancer’, ‘Small cell lung cancer’, ‘Pathways in cancer’ Suppl. File 3A – List of all enriched pathways). In skeletal muscle these genes are related to pathways matrix/ actin structure and remodelling, mechano-transduction, TGF-beta signalling and protein synthesis. Some of the same genes were included across these different KEGG cancer pathways, therefore, this related to 106 different cancer genes that were up or down regulated after acute RE across all 5 ‘cancer pathways’ (69 cancer genes were upregulated and 37 downregulated-see below for details). Pathway analysis on the methylome wide data also identified ‘Proteoglycans in cancer’ and ‘Pathways in cancer’ as significantly enriched (P ≤ 0.001), appearing within the top 20 KEGG pathways list after acute RE (Suppl. File 3B – List of all pathways). Comparison of the pathway diagrams demonstrated that there was a larger proportion of upregulated vs. downregulated genes included in ‘Pathways in Cancer’ and ‘Proteoglycans in cancer’ for the pooled transcriptome analysis (Suppl. Files 3C & 3D respectively). We also observed a larger number of hypomethylated vs. hypermethylated CpGs within ‘Pathways in Cancer’ and ‘Proteoglycans in cancer’ (Suppl. Files 3E & 3F respectively). Other than these two cancer pathways, none of the other top 20 enriched pathways in the pooled transcriptome gene expression analysis were the same as those identified in the top 20 enriched DNA methylation pathways in the methylome analysis (Suppl. File 3A and 3B, respectively).

Out of the genes up (69) and down regulated (37) across these ‘cancer’ pathways (Suppl. File 3G & 3H respectively), we then overlapped the genes that were hypo/hypermethylated respectively after acute RE in ^12^ (Figure 1C & D). This analysis identified 23 transcripts (Figure 1C; Suppl. File 3I) that were upregulated in the pooled transcriptome analysis that also demonstrated significant hypomethylation on the same gene following acute RE, this related to 29 CpG sites mapping to the 23 transcripts (as some genes had more than 1 CpG site modification per gene). This included genes: MSN, FOS, THBS1, ITPR3, TIMP3, RARA, FLNB, LAMA5, RASSF5, CRK, SMAD3, STAT3, COL4A1, ITGA2, WNT9A, ITGB3, KDR, ADCY3, CTTN, CD63, DOT1L, F2RL3 and GSK3B (Figure 1C; Suppl. File 3J). The largest proportion (57%) of the CpG’s were located in a promotor region for the same gene (for at least one their gene transcripts; Suppl. File 3J). In skeletal muscle most of these genes (13 out of 23) are associated with matrix / actin structure or remodelling and mechano-transduction (MSN, THBS1, TIMP3, FLNB, LAMA5, CRK, COL4A1, ITGA2, ITGB3, CD63, CTTN, RASSF5, F2RL3). With 3 genes associated with TGF-Beta signalling (SMAD3, FOS, WNT9A), 2 genes with calcium signalling (ITPR3, ADCY3), and 1 gene with IL-6 signalling (STAT3), protein synthesis (GSK3B) and retinoic acid signalling (RARA). Finally, after acute RE we identified 12 different genes (Figure 1D; Suppl. File 3K) that were downregulated in the comparative gene expression analysis that were also hypermethylated on 14 CpG sites in the methylome analysis ^12^, (as some genes had more than 1 CpG site modification). This included genes RUNX1T1, GAB1, ESR1, LAMA3, NANOG, SMO, ANK3, GADD45G, DROSHA, ATM, APAF1 and AGTR1 (Figure 1D; Suppl. File 3K), with varied functions in skeletal muscle (see discussion). Again, the largest proportion (64%) of the CpG‘s were also located in a promotor region for the same gene (for at least one of their gene transcripts; Suppl. File 3L).

### Pooled Transcriptome Analysis overlapped with the Methylome in Human skeletal Muscle after Chronic Resistance Exercise

A total of 181 biopsy samples made up the pooled chronic RE transcriptome analysis, with 86 (74 after outlier removal) baseline (pre) samples and 95 (87 after removal of outliers) post-chronic resistance exercise (RE) samples (See Suppl. Figure 2 and Suppl. File 1 for details). These studies shared 15,317 annotated genes by ‘gene symbol’ (Suppl. Figure 2A). Relevant QA/QC and outlier removal is depicted and described in Suppl. Figure 2B-R and the associated figure legend. From this pooled transcriptome analysis after chronic RE, we identified 2,448 genes that were significantly up (2,018 genes) or down (430) regulated (P ≤ 0.01) across all studies after chronic RE. Out of these 2,018 upregulated genes, 592 (Suppl. File 4B) were also hypomethylated after chronic RE (training/ termed loading in original article) from our methylome-wide analysis ^12^ (Figure 2A; Suppl. File 4B), compared with 516 genes that were upregulated and hypermethylated. Of the genes upregulated and hypomethylated, this equated to 793 different CpG sites mapping to these 592 genes (Suppl. File 4C). Furthermore, out of 430 downregulated genes, there were 98 genes across all transcriptome studies that were also hypermethylated in the methylome analysis (Figure 2B; Suppl. File. 4D), compared with similar number of genes (112) that were downregulated and hypomethylated. Of the genes downregulated and hypermethylated, this equated to 132 CpG sites on these 98 genes (Suppl. File 4E).

**Figure 2:**
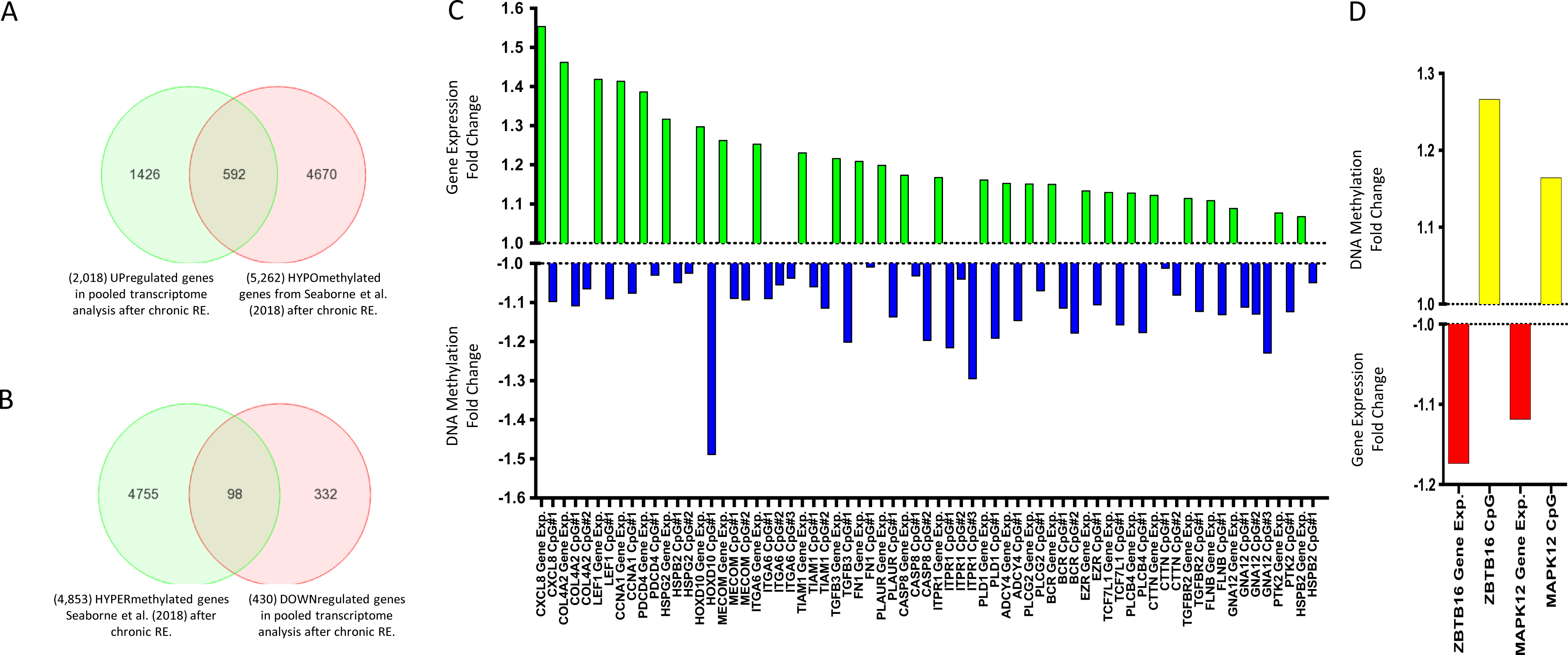
**A**, Venn Diagram analysis demonstrating out of 2,018 genes upregulated after chronic RE (p ≤ 0.01) in the pooled transcriptomic analysis, 592 of these genes were significantly hypomethylated in the methylome analysis after chronic RE^12^. **B**, Venn Diagram demonstrating out of 430 genes downregulated after chronic RE (p ≤ 0.01) in the pooled transcriptomic analysis, 98 of these genes were significantly hypermethylated in the methyome analysis after chronic RE ^12^. Note: 5262 / 4853 total hypo/hypermethylated CpGs after chronic RE is different than the original paper reporting 8,891 / 8,636 hypo/hypermethylated CpGs ^12^. This is due to the number of CpG’s that resided on the shared list of 15,317 annotated genes by ‘gene symbol’ across the pooled transcriptomic studies for chronic RE, depicted above in **5A**, in order to provide a direct comparison of CpG sites on the same genes from the pooled transcriptomic data set. **C**, ‘Cancer’ genes upregulated (GREEN bars) in pooled transcriptome studies (p ≤ 0.01) and hypomethylated (BLUE bars) in methylome analysis after chronic RE ^12^ (p ≤ 0.05). Fold change is Log transformed. In skeletal muscle, the majority of these genes are associated with matrix and actin structure / remodelling COL4A2, HSPG2, ITGA6, TIAM1, CTTN, GNA12, ADCY4, BCR, PLCG2, FN1, FLNB, PLAUR, EZR), mechano-transduction (PTK2/FAK and PLD1) and TGF-Beta signalling (TGFB3, TGFBR2, LEF1, MECOM). **D**, ‘Cancer’ genes downregulated (RED) in pooled transcriptomic studies and hypermethylated (YELLOW bars) in methylome analysis after chronic RE ^12^.

### Identification of significantly enriched KEGG pathways following chronic RE across the transcriptome and methylome

Analysis of the top 20 KEGG pathways significantly enriched in both the pooled transcriptome data sets and the methylome data set (Enrichment P values, ≤ 0.009 and ≤ 0.0001, respectively) following chronic RE, identified 7 pathways that were shared across both types of genome-wide analyses (Suppl. Files 5A and 5B, respectively). These 7 pathways, included: ‘Focal adhesion’, ‘circadian enrichment’, ‘glutamatergic synapse’ ‘phospholipase D signalling’, ‘human papillomavirus infection pathway’, ‘pathways in cancer’, and ‘proteoglycans in cancer’ pathways, and displayed a greater frequency of upregulated vs. downregulated gene transcripts and a higher number of hypomethylated vs. hypermethylated CpG sites following chronic RE (full gene lists and pathway diagrams for these pathways can be found in Suppl. Files 5C-P).

‘Pathways in cancer’ and ‘Proteoglycans in Cancer’ pathways after chronic RE were also identified in the above analysis to be significantly enriched after acute RE at both the transcriptome and methylome level. Furthermore, these were the only 2 out of 3 pathways that were in the top 20 enriched pathways identified in both acute and chronic RE pooled transcriptome analyses (the only other being the ‘Amoebiasis’ pathway). Therefore, as with the acute RE analysis above, we sought to analyse these cancer pathways in more detail after chronic RE. Indeed, these 2 cancer pathways combined included a total of 123 cancer related genes, 108 of which were unique genes (as some were shared/included in both cancer pathways). Out of these 108 genes that were up/down regulated we identified that 53 of these cancer genes were also modified at the DNA methylation level (Figure 2C). Of these 53 genes, this related to 91 different CpG’s (as some genes had more than 1 CpG site modification). From these 53 genes, 49 were upregulated (Suppl. File 6A) and we identified that 28 of these upregulated genes (Figure 2C; Suppl. File 6B) contained 40 hypomethylated CpG sites that mapped to these 28 transcripts, as some genes had more than modified CpG site (Figure 2C; Suppl. File 6C). This included genes: COL4A2, HSPG2, MECOM, TGFB3, ITGA6, TCF7L1, TIAM1, CTTN, CASP8, GNA12, ADCY4, BCR, PTK2, PLCG2, CCNA1, FN1, LEF1, PLCB4, ITPR1, PLD1, TGFBR2, PDCD4, FLNB, PLAUR, HOXD10, HSPB2, CXCL8 and EZR in muscle these genes are related to ECM/ actin structure and remodelling, mechano-transduction as well as focal adhesion (gene expression and methylation fold change of these genes can be seen in Figure 2C; Full lists in Suppl. File 6A & 6B). The largest majority (63%) of the CpG‘s were located in a promotor region for the same gene (for at least one of the genes transcripts) (Suppl. File 6C). In skeletal muscle, the predominant function for these genes, as with the acute RE analysis above, included ECM/ actin structure and remodelling, mechano-transduction (including focal adhesion) and TGF-Beta signalling. Furthermore, 4 out of the original 53 transcripts showed a significant down regulation in expression (Suppl. File 6D), with 2 of these transcripts (ZBTB16 and MAPK12) displaying a significantly hypermethylated CpG site within the promotor regions of the gene (for at least one of their known transcripts) following chronic RE (Figure 2D; Suppl. File 6E for gene expression and Suppl. File 6F for methylation).

### Epigenetically regulated muscle ‘memory’ genes highlighted in the analysis

It has been recently identified that following chronic RE training induced hypertrophy a number of CpG sites remained hypomethylated during detraining, even when exercise ceased and muscle returned to baseline/pre-exercise levels, with this hypomethylation also continuing through to a later period of retraining induced muscle hypertrophy ^12^. We therefore sought to identify if the same genes that were significantly regulated in the pooled transcriptome analysis after acute and chronic RE were also identified to be modified in the methylome after acute RE, training, detraining and retraining. Indeed, we identified 51 genes that were up/downregulated in both the acute and chronic RE transcriptome analysis, (full 51 gene list can be found in Suppl. File 7A&B for acute and chronic gene expression respectively), as well as significantly hypo/hypermethylated after acute RE (Suppl. File 7C), training/ loading (Suppl. File 7D), detraining/unloading (Suppl. File 7E) and retraining/reloading (Suppl. File 7F). All 51 genes were significantly differentially regulated and epigenetically modified in every condition on at least one of their CpG sites. However, because there was more than one CpG site per gene for several genes, there were only 13 of the same CpG sites (corresponding to 13 different gene transcripts) that were significantly modified across all methylation time points (acute RE, loading/training, unloading/detraining, retraining/reloading) and gene expression time points (acute/chronic RE). This included genes: FLNB, ASAP1, HMCN1, KDR, MBP, MYH9, MYH10, NDEL1, SRGAP1, SRGN, UBTF, ZMIZ1. From this we identified 5 CpG’s on 5 genes including: FLNB, MYH9, SRGAP1, SRGN, ZMIZ1 (with MYH9, SRGAP1, SRGN being promoter associated for at least one of their associated genes transcripts) with increased gene expression in both the transcriptome analysis after both acute and chronic RE and also demonstrated hypomethylation in these conditions (Figure 3 A respectively). Further analysis of the methylome-wide data set demonstrated retained hypomethylation in these CpG sites across detraining and retraining time points (Figure 3A). This retained epigenetic modification is most interesting following detraining (after training induced hypertrophy), when exercise was completely ceased and lean mass returned to baseline (pre-training) levels (reported previously ^12^), as this temporal regulation was demonstrated in ^12^ and proposed to identify those genes that have an epigenetic memory or prior exercise induced hypertrophy.

**Figure 3:**
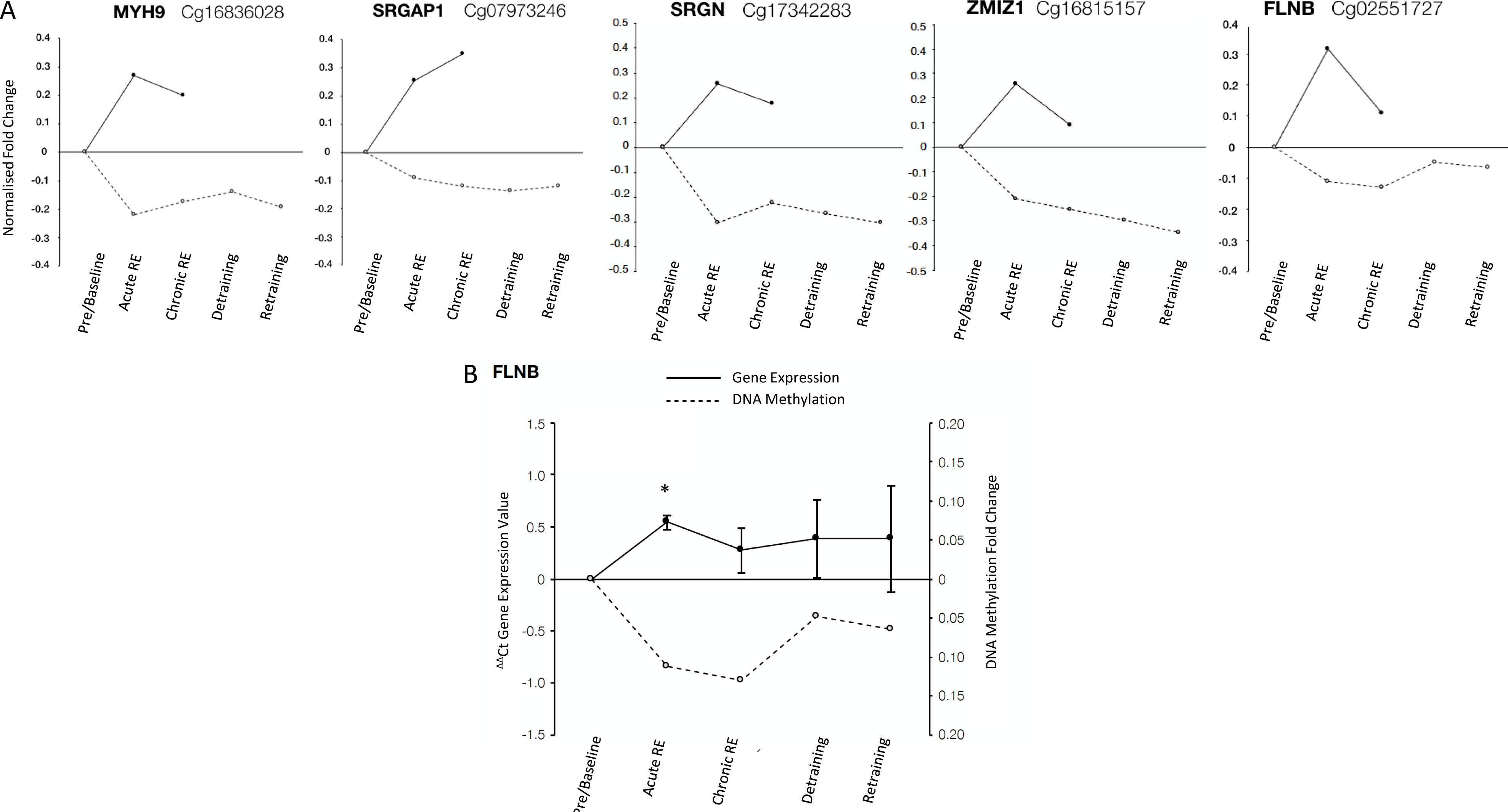
**A**, Fold change in gene expression (normalised to 0) from pooled transcriptome analysis after acute and chronic RE and methylome analysis after acute and chronic RE, detraining and retraining: MYH9, SRGAP1, SRGN, ZMIZ1 and FLNB (left to right) demonstrated increased gene expression in both the transcriptome analysis after both acute and chronic RE and also demonstrated hypomethylation in these conditions in the methylome analysis. Importantly, these genes also demonstrated retained hypomethylation even during detraining (following training induced hypertrophy) when exercise was completely cessed and lean mass in the original study ^12^ returned to baseline (pre-training) levels. **B**, New analysis of FLNB gene expression via rt-qRT-PCR (^ΔΔ^Ct gene expression value normalised to 0) demonstrated that the gene significantly increased in expression after acute RE vs. pre/baseline (P = 0.01). The average fold change in FLNB was also increased after chronic RE (training) and remained elevated vs. pre/baseline after detraining (unloading) and retraining (reloading) However, these increases failed to reach statistical significance. In the methylation array data FLNB methylation demonstrated an inverse association with the gene expression where the gene was hypomethlyated after acute and chronic RE, with sustained hypomethylation during detraining and retraining.

Finally, Filamin B (FLNB) was identified in the ‘cancer’ pathway (with a role in the actin cytoskeleton) in both acute RE and chronic RE transcriptomic analysis above, as well identified in the 51 genes significantly modified across transcriptome and methylome analysis, and further identified in the final 5 genes above with increased gene expression as well as hypomethylation (with retained methylation during detaining and retraining). Therefore, we undertook new analysis of the mRNA expression of this gene in samples derived from ^12^, after acute and chronic RE, detaining and retraining. Indeed, we identified that gene expression of FLNB was significantly increased after acute RE (1.54 fold ± 0.1 SEM vs. pre/baseline P = 0.01). The average fold change in FLNB was also increased after chronic RE (training) (1.27 fold ± 0.2 SEM vs. pre/baseline) and remained elevated vs. baseline after detraining (unloading) (1.38 fold ± 0.37 SEM vs. pre/baseline) and retraining (reloading) (1.38 fold ± 0.5 SEM vs. pre/baseline) (Figure 3B). However, these increases failed to reach statistical significance. Yet, this did demonstrate and inverse relationship with DNA methylation (hypomethylation after acute RE that was sustained throughout training, detraining and retraining, Figure 3B).

## Discussion

In the present study we undertook bioinformatic analysis and identified significantly up and down regulated genes across publicly available transcriptome studies after acute and chronic resistance exercise in human skeletal muscle. We then overlapped this with recent data that measured the DNA methylome response after acute resistance exercise (acute RE), chronic RE (loading/training), detraining (unloading) and retaining (reloading)^12^. We had the specific aim to identify novel genes/gene expression pathways that were epigenetically altered at the DNA methylation level after acute anabolic exercise, chronic resistance exercise induced hypertrophy and those associated with an epigenetic memory.

### KEGG Cancer Pathways are Significantly enriched in the Transcriptome and Methylome after Acute RE which are associated with ECM/ Actin Structure, Remodelling and Mechano-transduction in Human Skeletal Muscle

After acute RE we identified 866 up- and 936 down-regulated genes, with 270 (out of the 866 up-regulated) identified as being hypomethylated, and 216 (out of 936 downregulated) as hypermethylated. After KEGG pathway analysis, genes associated with several ‘cancer’ pathways were significantly enriched in both the pooled transcriptome analysis and also enriched in the methylome data after acute RE. This resulted in 23 upregulated and 12 downregulated ‘cancer’ genes that were also hypo/ hypermethylated, respectively. The largest proportion of the CpG sites (57%) on these ‘cancer’ associated genes were promoter associated. Modification of site specific methylation in the promotor region of coding gene sequences is known to be an important regulator of the down-stream expression of the transcript, largely due to manipulation of binding and accessibility of polymerase apparatus ^11^. We therefore identified an important role for DNA methylation in the promoter regions of genes within human skeletal muscle after acute RE, adding to previous work with similar findings in endurance exercise ^26^. Of these 35 ‘cancer’ genes, 23 were upregulated and hypomethylated, with a large proportion of these genes (13 out of 23) known to be important in matrix / actin structure and remodelling as well as mechano-transduction in skeletal muscle including: MSN, THBS1, TIMP3, FLNB, LAMA5, CRK, COL4A1, ITGA2, ITGB3, CD63, CTTN, RASSF5, F2RL3 (full gene names and general function is included in Suppl. File 8). Importantly, some of these ECM/actin structural and remodelling genes have been shown to demonstrate increases after aerobic or resistance exercise at the gene expression, protein abundance or activity level such as; TIMP3 ^27^, FLNB ^28^, COL4A1 ^29, 30^, or are involved in satellite/muscle cell regulation (ITGB3 ^31^, CTTN ^32^, LAMA5 ^33^, CRK ^34, 35^). However, to the authors knowledge there is no evidence to suggest that any of these genes are differentially methylated at the DNA level in skeletal muscle anabolism or hypertrophy. Therefore, the present study demonstrates for the first time, that important skeletal muscle genes/ gene pathways are also epigenetically regulated via DNA methylation, and that this predominantly occurs via the hypomethylation and upregulation of genes associated with structure and remodelling of the matrix and cytoskeleton as well as mechano-transduction in skeletal muscle. It is worth noting that a limitation of the acute RE analysis was that the acute pooled transcriptomic analysis included samples taken immediately post and up to 24 hrs after acute RE and that the overlapped methylome data was taken at 30 minutes post RE. This was due to few studies using the same timepoint making a pooled analysis difficult without adequate sample number. Furthermore, we have previously shown that DNA methylation changes at 30 minutes after acute RE can affect gene expression even after chronic RE training and retraining^12^. However, a time-course that includes multiple timepoints at both the transcriptomic and methylome level is required to directly investigate the temporal relationship between DNA methylation and gene expression after acute RE.

It is also worth noting that a further 10 genes after acute RE were significantly differentially regulated and enriched in ‘cancer’ pathways within both the transcriptome (upregulated) and methylome (hypomethylated) analysis, and included 3 genes associated with TGF-beta signalling (FOS, SMAD3, WNT9A), 2 genes with calcium signalling (ITPR, ADCY) and 1 gene with IL-6 signalling (STAT3), protein synthesis (GSK3B) and retinoic acid signalling (RARA). TGF-Beta signalling is a major growth regulator, and has been demonstrated to be important in skeletal muscle mass regulation and exercise adaptation. Indeed, we demonstrate increased expression and hypomethylation of FOS (or c-FOS) downstream of TGF-Beta, previously highlighted to be upregulated at the gene expression level after 30 minutes running ^36^ and after resistance exercise in rats ^37^, as well as post 2 hrs RE in humans ^38^. We also demonstrated increased expression/hypomethylation of SMAD3 (intracellular signal transducer and transcriptional modulator), known to be activated by TGF-beta and activin type 1 receptor kinases ^39^. However, in opposition with the current data, SMAD3 is increased in atrophy where muscle-specific knock-down of Smad2/3 in rodents protects from muscle atrophy following denervation ^40^. Yet, in support of the present study, it has recently been demonstrated that SMAD/JNK axis is important for muscle size regulation after resistance exercise, but not after aerobic exercise ^41^. Also, associated with TGF-Beta, the gene WNT9A was hypomethylated and upregulated, where it functions in the canonical Wnt/beta-catenin signalling pathway and has been identified, together with TGF-Beta2 and FGFR4, to be a key gene for the differentiation of muscle satellite cells ^42^. The calcium signalling genes that were upregulated and hypomethylated included; ITPR3 (or IP3R 3 - Inositol 1,4,5-trisphosphate receptor type/isoform 3) that has been associated with Ca2+ signals leading to gene regulation in skeletal muscle cells that contribute to muscle fibre growth and NMJ stabilization ^43^. While ADCY3 has no clearly defined role in skeletal muscle, it catalyzes the formation of the signalling molecule cAMP in response to G-protein signalling ^44^, and regulates Ca^2+^-dependent insulin secretion. Its methylation has been demonstrated to change in adipose tissue in response to altered nutrition ^45^, but no methylation data currently exists in skeletal muscle.

We also demonstrate increases in gene expression and hypomethylation of other important genes in muscle; STAT 3, GSK3B and RARA. Indeed, STAT3 is downstream of IL-6, and is known to play a critical role in regulating skeletal muscle mass, repair and is associated with myopathy (recently reviewed here ^46^). However, while it has been shown to have increased protein activity after resistance exercise ^47^, recent studies have shown that it is not required for overload induced hypertrophy in rodents ^48^. The important protein synthesis regulator, GSK3B (Glycogen synthase kinase-3 beta), regulates protein synthesis by controlling the activity of initiation factor 2B (EIF2BE/EIF2B5), and has previously been shown to increase after resistance exercise and decrease during skeletal muscle atrophy ^49^. Finally, RARA (Retinoic acid receptor alpha), important in skeletal muscle repair and satellite cell self-renewal ^49^ ^50^, is also upregulated and hypomethylated. Once again, to the authors knowledge there is no previous evidence to suggest that any of these transcripts are differentially methylated in skeletal muscle anabolism or hypertrophy. Further demonstrating the potential role DNA methylation plays in modulating the anabolic response of skeletal muscle mass after acute RE.

Of the above 35 ‘cancer’ genes up/downregulated after acute RE, we also demonstrated that 12 were downregulated and hypermethylated, including: RUNX1T1, GAB1, ESR1, LAMA3, NANOG, SMO, ANK3, GADD45G, DROSHA, ATM, APAF1 and AGTR1. These genes related to several different functions within skeletal muscle and with the exception of those relating to apoptosis (GADD45G, ATM, APAF1), display very limited connection to one-another. Most notably for skeletal muscle, however, ESR1 (estrogen receptor 1) in the present study was hypermethylated with reduced gene expression in the young adult males after acute RE, whereas in female mice after knock-down, it has been shown to cause muscle weakness ^51^ yet, overexpression can result in a slow muscle fibre phenotype ^52^. Therefore, a reduction observed in the adult males may therefore be associated with reduced slow fibre formation and subsequently may be conducive to faster fibre formation, previously identified to occur after resistance exercise. The association of hypermethylated and downregulation of ESR1 and its role in fibre type transition requires further investigation.

### KEGG Cancer Pathways are Significantly enriched after Chronic RE in the Transcriptome and Methylome and associated with ECM/ Actin Structure and Remodelling, Mechano-Transduction and Focal Adhesion in Human Skeletal Muscle

After chronic RE we identified 2,018 up- and 430 down-regulated genes with 592 (out of 2018 upregulated) identified as being hypomethylated, and 98 (out of 430 genes downregulated) as hypermethylated. As with the acute RE analysis above, KEGG pathway analysis identified both ‘Pathways in cancer’ and ‘Proteoglycans in cancer’ to be significantly enriched in both the pooled transcriptome data and methylome data. Furthermore, these were the 2 predominant pathways identified in both the acute and chronic transcriptome analysis. After chronic RE, 28 (out of 49 upregulated) and 2 (out of 4 down-regulated) ‘cancer’ genes were also hypomethylated/hypermethylated respectively. In conjunction with the acute RE data, the largest proportion (63%) of the CpG sites on these KEGG ‘cancer’ genes were promoter associated. This included: COL4A2, HSPG2, MECOM, TGFB3, ITGA6, TCF7L1, TIAM1, CTTN, CASP8, GNA12, ADCY4, BCR, PTK2, PLCG2, CCNA1, FN1, LEF1, PLCB4, ITPR1, PLD1, TGFBR2, PDCD4, FLNB, PLAUR, HOXD10, HSPB2, CXCL8 and EZR. Out of the 28 genes upregulated, a large proportion of the genes (14 out of 28) as with the acute RE analysis above, where associated with ECM/ actin structure and remodelling, mechano-transduction as well as focal adhesion in skeletal muscle including: COL4A2, HSPG2, TIAM1, CTTN, ADCY4, BCR, PTK2 aka. FAK, PLCG2, FN1, PLD-1, FLNB, EZR (for list of full gene name and broad function see Suppl. File 8). In terms of their role in skeletal muscle and exercise induced hypertrophy, only 7 out of these 14 upregulated and hypomethylated genes have been investigated, notably COL4A2, HSPG2, TIAM1, CTTN, PTK2/FAK, PLD1 and PLAUR. Indeed, the responsiveness of COL4A2 (Type IV collagen) to muscle unloading and reloading seems to be impaired in aged muscle of rats ^30^. Further, HSPG2 (Perlecan) has been identified to alter mass and metabolism in skeletal muscle ^53^. TIAM1 has been associated with skeletal muscle glucose uptake ^54^. CTTN (cortactin) contributes to the organization of the actin cytoskeleton ^55^ and is also important in actin filament remodelling in L6 myotubes ^32^. The abundance of FN1 (fibronectin) is blunted in elderly human skeletal muscle following muscle damage evoked via eccentric muscle contraction, compared to young healthy controls ^56^. PLAUR acts as a receptor for urokinase plasminogen (uPA) with uPA-null mice demonstrating a complete lack of hypertrophy in response to synergistic ablation ^57^. Also, while there are limited studies for the role of FLNB (filamin B) in skeletal muscle or resistance exercise induced hypertrophy, its loss has recently been highlighted to be associated with several myopathies and therefore a potential interesting target in skeletal muscle adaptation ^28^. Further, its closely related genes/protein family members, such as filamin A (FLNA) have been identified as a substrate of Akt and phosphorylated after endurance exercise, but not resistance exercise ^58^, and filamin C (FLNC) has been demonstrated to be closely associated with autophagy processes after resistance exercise ^59^. These findings therefore suggest FLNB requires further attention as to its role in skeletal muscle anabolism and hypertrophy.

Most interestingly in this group of identified hypomethylated and upregulated genes, PTK2 (Focal adhesion kinase/FAK) has been previously identified as an important regulator of mechanical loading in skeletal muscle, as it is predominantly located within the costamere (a mechano-sensory site of focal adhesion in the sarcolemma). FAK increases at the protein activity level after resistance exercise in human skeletal muscle, with its increased activation elevated to the largest extent after eccentric contraction ^60^. Furthermore, overexpression of FAK can have a protective effect on skeletal muscle reperfusion injury after limb ischemia ^61^, and increasing FAK can also increase the β1-integrin and meta-vinculin (+88%) content in skeletal muscle ^62^. PLD-1 is another major mechano-sensor that has been explicitly linked with numerous functions in skeletal muscle, where it has been shown to be fundamental for muscle cell differentiation ^63^, as well as myotube size via the activation of mechanistic target of rapamycin (mTOR) ^64^. Importantly in rodents, the activation of mTOR following eccentric lengthening contractions occurs through a PI3K-PKB-independent mechanism, a process that requires upstream PLD and phosphatidic acid ^65^.

The remaining genes in the 28 ‘cancer’ genes identified as upregulated and hypomethylated, as with the acute RE analysis, this included genes implicated in TGF-beta signalling (TGFB3, TGFBR2, LEF1 and MECOM) after chronic RE in human skeletal muscle. These associated transcripts included: TGFB3 (Transforming growth factor beta-3) itself, TGF-beta receptor type-2 (TGFBR2) a receptor for the TGF-beta ligands TGFB1, TGFB2 & TGFB3 as well as; LEF1, that is associated with WNT-TCF/LEF and TGF-Beta-SMAD3 signalling in controlling muscle stem cell proliferation and self-renewal ^66^. Finally, MECOM (alias EVI-1) has been demonstrated to play a role in growth regulation via TGF-beta/SMAD signalling^67^. However, to the authors knowledge, MECOM has not been studied specifically in skeletal muscle anabolism or hypertrophy.

Collectively, we identify important epigenetically regulated ‘cancer’ genes that are associated with ECM/Actin structure and remodelling, mechano-transduction/focal adhesion (e.g. FAK and PLD) and TGF-Beta signalling after chronic RE in human skeletal muscle. While these were also the gene pathways associated with acute RE analysis above, it is worth noting that only two of the same genes, CTTN and FLNB were increased and hypomethylated after both acute and chronic resistance exercise analysis, and there were no other genes that were downregulated and hypermethylated that appeared in both acute and chronic transcriptome analysis. This suggested that while ‘cancer’ pathways, important in skeletal muscle matrix/ actin structure and remodelling, mechano-transduction and TGF-Beta signalling were important after both acute and chronic resistance exercise, the longer-term adaption to chronic exercise seems to be regulated by different genes within the same pathways versus a single bout of resistance exercise.

### Newly Identified Epigenetically Modified ‘Memory’ Genes in Human Skeletal Muscle

51 genes were identified to be up/downregulated in both the acute and chronic RE transcriptomic analysis as well as significantly hypo/hypermethylated after acute RE, chronic RE, detraining and retraining. Five genes (MYH9, SRGAP1, SRGN, ZMIZ1, FLNB) demonstrated increased gene expression in the acute and chronic RE transcriptome while also demonstrating hypomethylation in these conditions. Importantly, these 5 genes also retained hypomethylation even during detraining (following training induced hypertrophy) when exercise completely cessed and lean mass returned to baseline (pre-training) levels. We therefore suggest, given the retention of epigenetic information within CpG sites of these transcripts, that these may be newly identified memory genes that warrant further investigation. Interestingly MYH9, while it has been shown to be expressed at the gene level in skeletal muscle tissue, it is not thought to be translated into the protein and is therefore classed as a non-muscle myosin heavy chain. Therefore, it could be hypothesised that its hypomethylation and increased gene expression has other, yet undescribed, post-transcriptional functions in skeletal muscle anabolism and hypertrophy. SRGAP1 is a GTPase-activating protein for RhoA and Cdc42 small GTPases ^68, 69^ and therefore, while potentially important in mechano-transduction, has not been studied in skeletal muscle. Serglycin (SRGN) is a proteoglycan that has been shown to increase its expression after acute and chronic aerobic exercise in humans, with cultured human muscle derived cells both expressing and secreting serglycin^70^. However, to the authors knowledge SRGN has not been studied after resistance exercise at either the gene expression of DNA methylation level, yet we identify it as a potential gene involved in epigenetic muscle memory. ZMIZ1 is a transcriptional co-activator that modules the transcription of androgen receptor (AR), SMAD3/4, and p53 in other cell types ^71–73^. However, its role in skeletal muscle is yet undefined. Intriguingly, AR, SMAD and p53 are all important regulators of skeletal muscle mass and metabolism and therefore this gene requires further investigation as to its epigenetic role in skeletal muscle.

Finally, Filamin B (FLNB) was also identified in the gene expression and methylation analysis in both acute and chronic pooled transcriptome data and was significantly enriched in ‘cancer pathways’ in both the pooled transcriptome and methylome. At the protein level filamin B is involved in connecting the cell membrane constituents to the actin cytoskeleton, where mutations of the gene are inextricably linked to several myopathies ^28^. From an exercise-induced response/adaptation perspective, there is limited information regarding the role of Filamin B in skeletal muscle following RE. Interestingly, Filamin A (protein family member) has been shown to be a substrate of Akt, and to be phosphorylated after endurance exercise, but not after resistance exercise ^58^. More so, Filamin C has been shown to be altered following RE, and is closely associated with autophagy processes altered after resistance exercise ^59^. Experimentally, we were able to confirm that Filamin B was increased at the gene expression level after acute and chronic RE and remained elevated (but non-significantly) after detraining and retraining where the gene remained as hypomethylated even during exercise cessation (detraining). However, increases in FLNB were only significant after acute RE. Despite this, and given the associated sustained hypomethylation, we suggest that this gene requires further mechanistic investigation as to its role in skeletal muscle anabolism, hypertrophy and memory.

## Conclusion

Importantly, for the first time across both the transcriptome and methylome, this study identifies novel epigenetically modified genes that are associated with gene expression changes after acute and chronic resistance exercise and muscle memory. Those identified suggest predominant roles for genes associated with human skeletal muscle ECM/ actin structure and remodelling, mechano-transduction as well as TGF-beta signalling associated with anabolism, hypertrophy and memory.

## Acknowledgments

Genome-wide methylation studies were funded by a GlaxoSmithKline grant awarded to Adam P. Sharples (PI). Daniel Turner’s PhD is supported by the Society for Endocrinology, UK.

## Author Contributions

Sharples, Turner and Seaborne, conceived and designed the research, analysed all data, performed the research and wrote the manuscript. All authors reviewed the manuscript drafts and inputted corrections, amendments and their expertise.

---

**Suppl. Figure 1: A**, Venn diagram analysis identified a shared list of 14,992 annotated genes by ‘gene symbol’ across the pooled transcriptomic studies for acute RE. Note ^4, 5^ use the same gene array platform but the number of genes is depicted as different. This is due to the highest probe set analysis (see methods) altering the genes that could be compared in this study. Inspecting **B**, box and whisker plots before batch correction and, **C**, frequency plots by lines there was a noticeable signal/ value variation as a result of batch effects across studies. **D**, Box and Whisker plots post batch correction and **E**, frequency normalisation plots by lines post batch correction**. F**, PCA post batch correction by time (pre/post) and **G:** PCA by study depicting the outliers removed (line strikethrough) that were located outside 2SD’s of the centroid value using ellipsoids. **H**, Sample box and whiskers plot with batch correction and outlier samples (highlighted in F/G) removed. **I**, Frequency Plot by Lines after batch correction and with outlier samples (highlighted in F/E) removed.

**Suppl. Figure 2: A**, Venn diagram analysis identified a shared list of 15,317 annotated genes by ‘gene symbol’ across the pooled transcriptomic studies for chronic RE. **B**, PCA for study and **C**, time (pre / post) **D**, box and whisker plots by study and **E**, by time point. **F**, frequency/density plots by lines for study and **G** time (pre/post). There was noticeable signal/ value variation as a result of batch effects across studies. Following batch correction: **H**, PCA by time and **I**, by study demonstrated smaller variation than prior to batch correction (B & C above). **J** Box and whisker plots by study and **K**, by time post batch removal. **L**, Frequency plots by time and **M**, by study post batch removal demonstrated more appropriate signals and distribution compared with those prior to batch removal (D-G). **N**, PCA post batch correction by time (pre/post) depicting the outliers removed (line strikethrough) that were located outside 2SD’s of the centroid value using ellipsoids. **O**, Frequency/density plots by time and **P**, by study post batch removal with outliers (identified in N) removed. **Q**, Box and whisker plots by time and **R**, by study post batch removal with outliers (identified in N) removed.

