## Supplementary material for "Comparative Transcriptome and Methylome Analysis in Human Skeletal Muscle Anabolism, Hypertrophy and Epigenetic Memory": Suppl File 1

United Kingdom

**Table 1**. **Gene expression transcriptome studies pooled for analysis of gene expression for both acute and chronic resistance exercise (RE)**. Note: There were only 2 studies that contained relevant rested baseline samples prior to (pre) chronic RE within the same study, severely limiting the number of pre/rested samples for the chronic RE analysis [5,9], whereas a total of 4 studies had rested post chronic resistance exercise samples ([5, 8, 9, 10] with [8,10] without relevant resting baseline/pre samples). We therefore included the samples from the pre-acute RE transcriptome analysis from the three studies of [3,4,6] as relevant pre/baseline rested samples.

| Acute RE Studies | PMID | GEO Acc. | Array Platform | Exercise | Time point | Notes |
|---|---|---|---|---|---|---|
| MacNeil et al. (2010)[3] | 20502695 | GSE19062 | GPL6255 Illumina humanRef-8 v2.0 | Eccentric exercise of the quadriceps, 15 sets of 10 reps. maximally resisting flexion of the knee at 120°/s, 1 min rest between sets. | Pre, 3h Post | E2 supplement group was removed. Only placebo/no supplement group analysed. |
| Raue et al. (2012)[5] | 22302958 | GSE28422 | GPL570 [HG U133_Plus_2] | Bilateral knees extension of the quadriceps, 3 sets of 10 reps at 70-75% of their 1RM | Pre, 4h Post | Elderly male/ female and young female adult acute and chronic RE groups were removed. Only young male adult group analysed. |
| Vissing & Schjerling. (2014)[6] | 25984345 | GSE59088 | GPL6244 [HuGene-1_0-st] | 3 separate exercises of the quadriceps x 1 set of their 12 RM, 1.5 mins. rest between exercises. | Pre, 2.5h, 5h Post | Endurance exercise group was removed. Only pre and post-acute RE were analysed. |
| Murton et al. (2014)[4] | 24265280 | GSE45426 | GPL570 [HG U133_Plus_2] | Knee extension, 5 sets of 30 maximal isokinetic contractions at 180°/s, 1 min rest between sets. | Pre, 24h Post | Non-exercise group was removed. Pre/post-acute RE group used for the analysis. |
| Lundberg et al. (2016)[7] | 27101291 | GSE74194 | GPL17692 [HuGene-2_1-st] | Knee extension, 4 sets of 7 reps (70% of max.), 2 min rest between sets. | Post 3h Only | No relevant pre, as the comparison was post-acute RE from one limb (that performed RE) vs. the contralateral limb that underwent both RE plus endurance exercise. Post RE limb samples used for the analysis only. RE+ endurance limb samples were removed. |
| **Chronic RE Studies** | | | | | | |
| MacNeil et al. (2010)[3] | 20502695 | GSE19062 | GPL6255 Illumina humanRef-8 v2.0 | Pre (rested) samples used in analysis only. | Pre Only | Pre (resting) samples (from acute RE transcriptome analysis above) only were included in the analysis. E2 supplement group was removed. Only placebo/no supplement group analysed. |
| Liu et al. (2010)[8] | 21106073 | GSE24235 | GPL570 [HG U133_Plus_2] | 2/wk by 12 wks progressive RE of the arms (5 elbow flexor exercise x 3 sets of 6 RM). | Post Only | Analysed the resting samples only from biopsies of the arm that had done the 12 weeks chronic RE. |
| Raue et al. (2012)[5] | 22302958 | GSE28422 | GPL570 [HG U133_Plus_2] | 3/wk by 12 wks Bilateral knees extension of the quadriceps, 3 sets of 10 reps at 70-75% of their 1RM. | Pre, Post | Elderly male / female and young female adult acute RE groups were removed. Only young male adult group pre and post chronic RE at rest were analysed. |
| Phillips et al. (2013)[9] | 23555298 | GSE47881 | GPL570 [HG U133_Plus_2] | 3/wk by 20 wks progressive RE (4 wks 40-60% 1RM, 16 wks 70% 1RM) 1RM assessed every 4 wks. Multiple sets (no. not described) x 12 reps. | Pre, Post | Resting Pre and Post Chronic samples analysed. |
| Thalacker-Mercer et al. (2013)[10] | 23632419 | GSE42507 | GPL6480 Agilent-014850 | 3/wk by 16 wks RE (specific intensity or exercises performed not described) | Post Only | Non-responder group was removed. Moderate and extreme responders analysed only. |
| Vissing & Schjerling. (2014)6 | 25984345 | GSE59088 | GPL6244 [HuGene-1_0-st] | Pre (rested) samples used in analysis only. | Pre Only | Pre (resting) samples (from acute RE transcriptome analysis above) were included in the analysis. |
| Murton et al. (2014)[4] | 24265280 | GSE45426 | GPL570 [HG U133_Plus_2] | Pre (rested) samples used in analysis only. | Pre-Only | Pre (resting) samples (from acute RE transcriptome analysis above) were included in the analysis. |
