## Supplementary material for "Comparative Transcriptome and Methylome Analysis in Human Skeletal Muscle Anabolism, Hypertrophy and Epigenetic Memory": Supple File 8

United Kingdom

**Suppl. File 8:**
**13 genes upregulated and hypomethylated after acute RE involved in ECM/Actin structure/remodelling and mechanotransduction.**

**MSN** (Meosin) involved in actin and plasma membrane crosslinking and associated with muscle dystrophies [1].

**THBS1** (Thrombospondin 1) an adhesive glycoprotein mediating cell-to-cell and matrix interactions that can bind to fibrinogen and as well as ECM proteins such as laminin, collagen and integrins, with a demonstrated role in capillary density in muscle [2].

**TIMP3** (Metalloproteinase inhibitor 3) involved in remodelling the matrix [3].

**FLNB** (Filamin B) that connects the cell membrane to actin cytoskeleton, and mutations that lead to myopathies [4].

**LAMA5** (Laminin 5) an ECM protein that is altered after remodelling of the muscle matrix [5];

**CRK** (Adapter molecule crk) regulates cell adhesion associated with mechano-transduction via FAK [6,7].

**COL4A1** (Collagen alpha-1 IV chain) is a major structural component of basement membranes, linking to other laminins and proteoglycans[8].

**ITGA2** (Integrin alpha-2/beta-1) is a receptor for laminin, collagen, fibronectin and E-cadherin.

**ITGB3** (Integrin Beta-3) together with Integrin Alpha 5 is a receptor for ECM proteins in muscle, and important in muscle cell migration [9].

**CD63** (CD63 antigen), functions as cell surface receptor for TIMP1[10] and is involved in the activation of integrin/FAK/AKT signalling.

**CTTN** (Src substrate cortactin) contributes to the organization of the actin cytoskeleton [11] and actin filament remodelling in L6 myotubes [12].

**F2RL3** (aka PAR-4 Proteinase-activated receptor 4) is a receptor for activated thrombin or trypsin, however it demonstrated no real role the in differentiation in muscle cells [13].

**RASSF5** (Ras association domain-containing protein 5 aka. RAPL), although no known role in skeletal muscle, together with RAP1A is involved in extension of microtubules in endothelial cells [14].

**<u>14 genes upregulated and hypomethylated after chronic RE involved in ECM/Actin structure/remodelling and mechanotransduction.</u>**

**COL4A2** (Type IV collagen) structural protein in the ECM.

**HSPG2** aka. Perlecan (Basement membrane-specific heparan sulfate proteoglycan core protein). Important component of basement membranes

**ITGA6** (Integrin alpha-6) is a receptor for laminin. ITGA6:ITGB4 binds to IGF-I/2 and this binding is essential for IGF-I/2 signaling.

**TIAM1** (**T-lymphoma invasion and metastasis-inducing protein 1)** that connects extracellular signals to cytoskeletal activities.

**CTTN** (cortactin) contributes to the organization of the actin cytoskeleton.

**GNA12** (Guanine nucleotide-binding proteins-G proteins) involved as transducer in transmembrane signalling.

**ADCY4** (Adenylate cyclase type 4) involved in G-protein signalling.

**BCR** (Breakpoint cluster region protein) is a GTPase-activating protein.

**PTK2 aka. FAK** (Focal adhesion kinase) located in the mechano-sensing costamere of skeletal muscle; PLCG2, an enzyme involved in transmembrane signalling.

**FN1** (Fibronectin) binds cell surfaces including collagen, fibrin, heparin, and actin; PLD-1; (Phospholipase D1) involved in transmembrane trafficking.

**FLNB** (Filamin-B) that connects cell membrane constituents to the actin cytoskeleton.

**PLAUR** (Urokinase plasminogen activator surface receptor) acts as a receptor for urokinase plasminogen activator.

**EZR** (Ezrin) involved in connections of major cytoskeletal structures to the plasma membrane.
